# Development of a novel *Francisella tularensis* Live Vaccine Strain expressing ovalbumin provides insight into *Francisella tularensis*-specific CD8^+^ T cell responses

**DOI:** 10.1101/200592

**Authors:** David E. Place, David R. Williamson, Yevgeniy Yuzefpolskiy, Bhuvana Katkere, Surojit Sarkar, Vandana Kalia, Girish S. Kirimanjeswara

## Abstract

Progress towards a safe and effective vaccine for the prevention of tularemia has been hindered by a lack of knowledge regarding the correlates of protective adaptive immunity and a lack of tools to generate this knowledge. CD8^+^ T cells are essential for protective immunity against virulent strains of *Francisella tularensis*, but to-date, it has not been possible to study these cells in a pathogen-specific manner. Here, we report the development of a tool for expression of the model antigen ovalbumin (OVA) in *F. tularensis*, which allows for the study of CD8^+^ T cell responses to the bacterium. We demonstrate that in response to intranasal infection with the *F. tularensis* Live Vaccine Strain, pathogen-specific CD8^+^ T cells expand after the first week and produce IFN-γ but not IL-17. Effector and central memory subsets develop with disparate kinetics in the lungs, draining lymph node and spleen. Notably, *F. tularensis*-specific cells are poorly retained in the lungs after clearance of infection. We also show that intranasal vaccination leads to more pathogen-specific CD8^+^ T cells in the lung-draining lymph node compared to scarification vaccination, but that an intranasal booster overcomes this difference. Together, our data show that this novel tool can be used to study multiple aspects of the CD8^+^ T cell response to *F. tularensis*. Use of this tool will enhance our understanding of immunity to this deadly pathogen.

## INTRODUCTION

The Gram-negative bacterium, *Francisella tularensis*, is the etiological agent of the disease tularemia in humans. Inhalation of approximately 10-50 CFU [1] of *F. tularensis* subsp. *tularensis* can lead to severe and rapidly-progressing disease, which is associated with high mortality without early intervention [2]. Additionally, the bacterium is easily aerosolized [3], and can be genetically manipulated to render it antibiotic resistant. The combination of these factors makes *F. tularensis* an ideal candidate biological weapon. Indeed, it was developed for this purpose by several countries in the 20^th^ century [2,4–6], and remains a tier 1 select agent due to the potential for use as an agent of bio-terrorism.

There is currently no approved vaccine for the prevention of tularemia. An empirically attenuated Live Vaccine Strain (LVS), derived from a *F. tularensis* subsp. *holarctica* isolate, was developed over 50 years ago [7]. The exact basis of attenuation, however, is still not well defined; this and the potential for either loss of protectiveness [8,9] or reversion to virulence [10,11] are barriers for the approval of LVS for vaccination in humans. Additionally, the effectiveness of LVS in generating long term protection from respiratory challenge with virulent strains is poor in many models [12–14]. To facilitate the development and approval of a vaccine that is safe and effective, it is crucial that the correlates of protective adaptive immunity to *F. tularensis* be clearly defined.

Antibody-mediated immunity appears to be a poor correlate of immunity to highly virulent *F. tularensis* strains; antibody titers do not correlate with protection in humans[15], and the transfer of immune serum fails to protect recipient mice against the challenge with virulent strain of *F. tularensis* [16–18]. In contrast, both CD4^+^ and CD8^+^ T cells are known to be required for protection, as depletion of either subset abolishes protective immunity [12,19,20]. To truly hone in on correlates of protective T cell responses, it is necessary to be able to differentiate cells specifically responding to the pathogen of interest from cells of other specificities [21]. These non-specific cells may be far more abundant than pathogen-specific cells, thus representing a significant level of ‘background noise’ that may mask important insights into the true response to the pathogen. Antigen-specific cells can be studied by staining with MHC-peptide tetramers [22], or by tracking adoptively transferred transgenic T cells that are specific for a pathogen epitope. Thus far, there has been no success in using MHC-peptide tetramers to track T cells specific to natural *F. tularensis* antigens and no TCR-transgenic mice that recognize Francisella-specific epitopes have been generated. An alternative approach involves engineering strains of *F. tularensis* that express model antigens, which can be studied using existing tools.

In this regard, Roberts et al. have developed a construct in which they express the *F. tularensis* protein IglC tagged with the gp61-80 epitope of LCMV, allowing for tracking of antigen-specific CD4^+^ T cell responses using MHC-II tetramers [20]. This tool has allowed investigators to characterize antigen-specific CD4^+^ T cells in various contexts and begin identifying the correlates of CD4-mediated protection from tularemia. For instance, a protective vaccine leads to more antigen-specific CD4^+^ T_EM_ in the mediastinal lymph node (MLN) and spleen, as compared to a non-protective vaccine [20]. Additionally, the tool has been used to study how these cells respond to a prime-boost strategy [13] and has revealed the dramatic influence high avidity CD4^+^ T cell epitopes have on protection [13,20]. While this tool will undoubtedly yield many more insights into the role of CD4^+^ T cells in immunity to *F. tularensis,* no such tool currently exists to characterize CD8^+^ T cells, which are also required for protection against this predominantly intracellular pathogen [12,19,20].

To address this shortcoming, we have developed a tool for expression of chicken ovalbumin (OVA) protein in *F. tularensis*. This tool makes it possible to study the response of OT-I (TCR-transgenic) CD8^+^ T cells [23], which can be adoptively transferred into congenic C57BL/6 mice. Here we report the development of this tool and a proof-of-concept study using OVA-expressing *F. tularensis* LVS (termed LVS-OVA). In response to LVS-OVA, OT-I CD8^+^ T cells proliferate, differentiate into effector and central memory subsets, and produce interferon gamma (IFN-γ). We also compare how these cells respond following intranasal or scarification vaccination with LVS-OVA, followed by an intranasal booster. This novel tool will enable further detailed studies into the CD8^+^ T cell response to *F. tularensis*.

## MATERIALS AND METHODS

### Mouse Strains

C57BL/6J (Thy1.2^+^), C57BL/6J Thy1.1^+^ congenic mice (B6.PL-Thy1^a^/CyJ), and OT-I (C57BL/6-Tg(TcraTcrb)1100Mjb/J), mice were purchased from The Jackson Laboratories (Bar Harbor, ME). Unless otherwise indicated, experiments were performed using heterozygous wild-type Thy1.1^+^/Thy1.2^+^ recipient mice with OT-I (Thy1.1−/Thy1.2^+^) donor splenocytes. All mice were maintained in specific pathogen-free conditions at the Pennsylvania State University animal care facilities.

### Bacterial Strains and Cell Culture

Cloning was performed using *Escherichia coli* DH5α grown at 37°C in LB broth or LB agar containing tetracycline or kanamycin (10μg/mL or 50μg/mL, respectively). *Francisella tularensis* Live Vaccine Strain (LVS), obtained from Albany Medical College, was grown shaking in BHI broth at 37°C or on modified Mueller-Hinton II agar supplemented with hemoglobin (Thermo Scientific) and IsoVitalex (BD Biosciences) at 37°C in 5% CO_2_ with tetracycline or kanamycin (2μg/mL or 10μg/mL, respectively). For allelic exchange, Km^R^ LVS primary recombinants were streaked out on plates containing 5% sucrose and resulting colonies were screened for secondary recombination by plating single colonies on antibiotic-free, ampicillin, or kanamycin plates as previously described [24]. Mid-log phase liquid cultures of LVS were frozen at −80°C directly in BHI and enumerated regularly to calculate doses for mouse inoculation.

For immunoblots, liquid cultures (OD_600_ ~0.10) were pelleted and resuspended in bacterial lysis buffer followed by sonication. Lysate was mixed with 5X Laemelli buffer (BioRad) before running on 12% pre-cast Criterion TGX gels (BioRad). Protein was transferred to PVDF membrane using the Trans-Blot Turbo mixed molecular weight program. Membranes were then blocked in 5% BSA/TBS-T for 1h at room temperature, probed with anti-6xHis-Biotin (1:5000, Rockland #600-406-382) in 5%BSA/TBS-T for 1h at room temperature, washed, and probed with a secondary streptavidin-HRP (1:50000, Jackson ImmunoResearch) for 1h at room temperature. Blots were developed using Millipore Immobilon Chemiluminescent Substrate and images were collected on a BioRad ChemiDoc XRS+.

### Plasmids

Plasmid pKK214 containing GFP was obtained from Dr. Karl Klose and the GFP gene was removed by restriction digestion and replaced with a cloning site containing XbaI restriction sites. LVS genomic DNA was used to amplify the promoter for the bacterioferritin gene *(bfrp)* by PCR and to introduce flanking XbaI and BamHI restriction sites. The complete *vgrG* gene was amplified with primers introducing a BamHI site and replacing the stop codon with an XhoI site. To generate the codon-optimized ovalbumin epitope tag (containing OVA amino acids 239-345, a 6xHis tag, stop codon, and flanking XhoI and XbaI sites), the LVS codon frequency table from the Codon Usage DB (http://www.kazusa.or.jp/codon/cgi-bin/showcodon.cgi?species=376619) was imported into OPTIMIZER (genomes.urv.es/OPTIMIZER) using the standard setting of “one amino acid - one codon” [26]. The codon-optimized sequence was submitted for synthesis to Integrated DNA Technologies as a gBlock Gene Fragment, cloned into pCR4 cloning vector (Invitrogen) and confirmed by sequencing by the Penn State Genomics Core Facility. The bacterioferritin promoter, vgrG, and OVA_239-345_-6xHis fragments were ligated with T4 DNA ligase (NEB) and cloned into Xbal-digested pKK214 to generate pKK214-vgrG-OVA or pKK214-OVAgB. Electroporation was used to introduce plasmids to *E. coli* and LVS. A summary of primers, plasmids, and strains used can be found in Table 1.

**Table 1:**
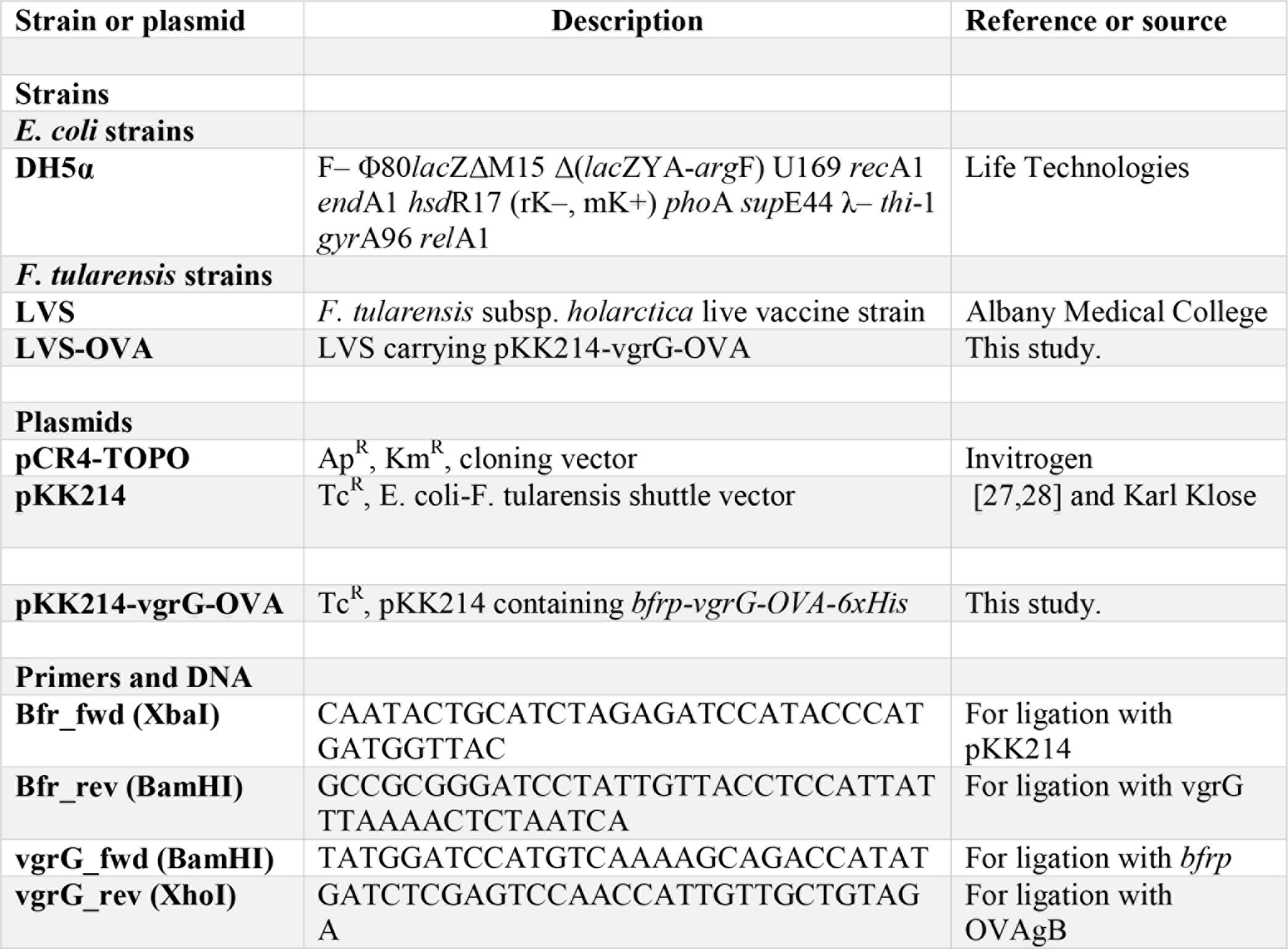
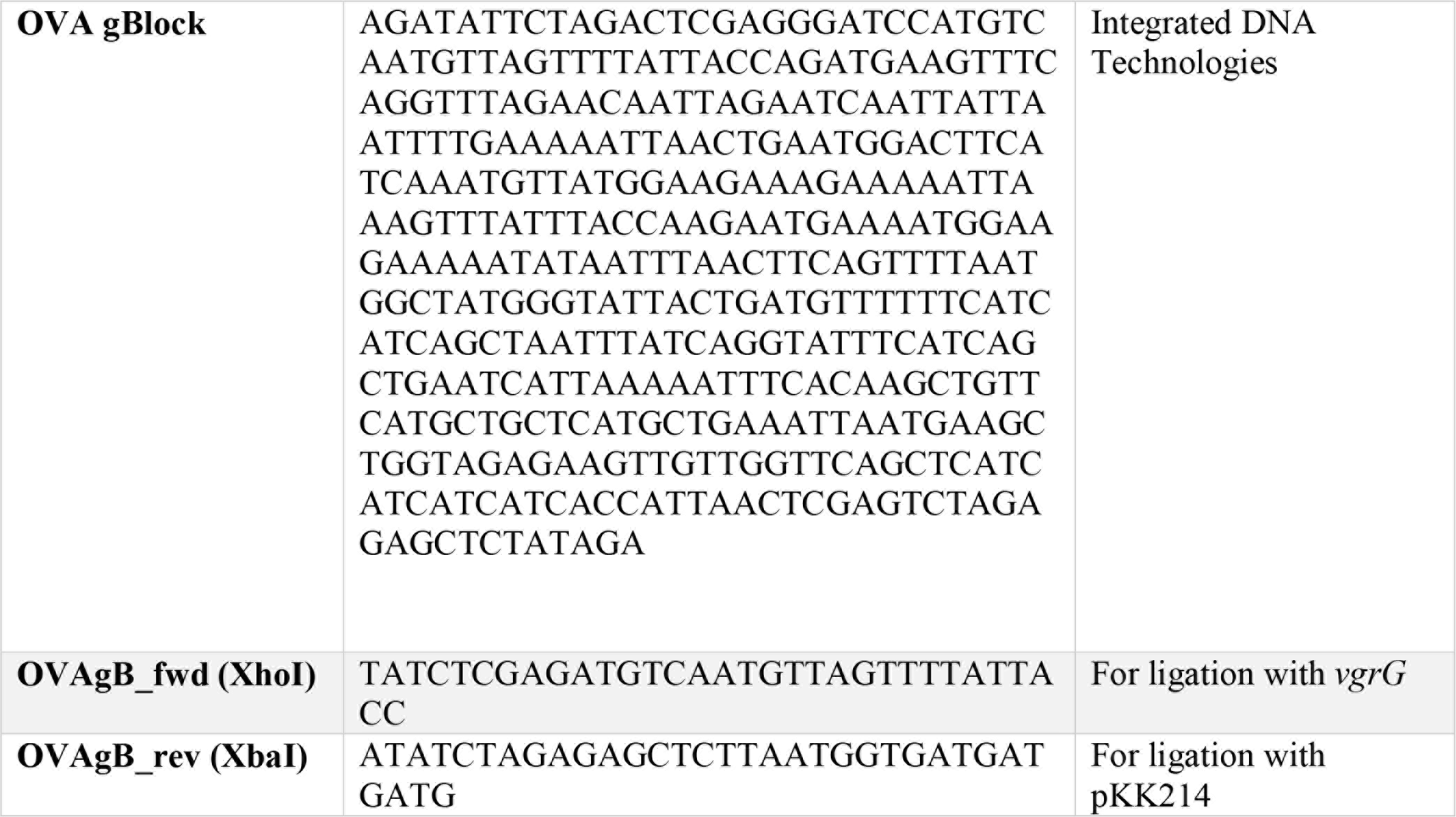
a summary of primers, plasmids and bacterial strains used in this study.

### Adoptive Transfer

For adoptive transfer of OT-I splenocytes (Thy1.1^−^/Thy1.2^+^), sex-matched spleens were aseptically collected in RPMI 1640 containing 10% FBS, 2mM L-glutamine, 1mM sodium pyruvate, 1x non-essential amino acids, 10mM HEPES, 55μM beta-mercaptoethanol and penicillin/streptomycin. Spleens were passed through a 70μm mesh filter by crushing with a 3mL syringe plunger and pelleted. Red blood cells were lysed with sterile 0.84% ammonium chloride/EDTA/Sodium bicarbonate and washed with complete media. For experiments with CFSE-labelled splenocytes, cells were washed with plain RPMI 1640 before resuspension at a concentration of 10^7^ cells/mL in RPMI 1640 containing 5μM CFSE. Staining was quenched by addition of 10% FBS and cells were washed twice with complete RPMI. 5×10^6^ OT-I splenocytes were transferred to Thy1.1^+^/Thy1.2^+^ recipients by intraperitoneal injection 24h before bacterial inoculation.

### Mouse Infection

Mouse inoculation was performed by thawing enumerated frozen bacterial stocks and serially diluting in PBS to the desired dose, which was 1000 CFU for standard intranasal experiments, 10,000 CFU for intranasal boosters, and 100,000 CFU for skin scarification. For intranasal inoculation, mice were lightly anesthetized using isoflurane and a 50μL droplet was administered to the external nares and inhaled. For skin scarification, mice were anesthetized using isoflurane, the fur of the upper back was shaved, and the skin cleaned by wiping with 70% ethanol. After drying, a 10μL droplet was administered to the bare skin, and a sterile bifurcated needle (Precision Medical Products) used to prick the skin 15 times. Mice were euthanized by CO_2_ inhalation and dissected aseptically in a biosafety cabinet. For experiments including bacterial burden determination, serial dilutions of single cell suspensions were prepared as described below, and plated on modified Mueller-Hinton agar plates.

### Flow Cytometry and Intracellular Cytokine Staining

Mice were euthanized and cold PBS was used to perfuse tissues by cardiac injection. The caudal mediastinal lymph node (MLN) and spleen were collected in HBSS with 1.3mM EDTA and crushed with a 3mL syringe plunger through a 70μm nylon filter. Lungs were collected, minced, suspended in HBSS with 1.3mM EDTA, and shaken 30 min. at 37°C. Following this, lungs were pelleted, resuspended in complete RPMI containing 200U/mL collagenase type I (Worthington Biochemical Corporation) and shaken at 37°C for 1h before passing through a 70μm nylon filter. Spleen and lung cells were treated with RBC lysis solution and washed with PBS/2%FBS. For surface staining, cells were incubated with relevant antibodies for 30 min on ice, washed, and fixed in PBS/1% PFA prior to running on a BD LSR Fortessa at Penn State Microscopy and Cytometry Facility. Intracellular cytokines were analyzed by stimulating ~2-10×10^6^ cells in 96-well plates at 37°C and 5% CO_2_ in the presence of 1xBrefeldin A (BioLegend) alone (unstimulated), or with 8nM phorbol 12-myristate 13-acetate (PMA) (Sigma) / 500ng/mL Calcium Ionophore A23187 (Sigma #C7522) for 4h in complete RPMI. Cells were stained for extracellular markers as above, fixed, then permeabilized with 0.1% saponin in PBS/2%FBS. Antibodies for intracellular cytokines were incubated 30min in permeabilization buffer, washed 3x with permeabilization buffer then 3x with PBS/2%FBS, and analyzed immediately. Analysis was performed using FlowJo v10.0.8 software (company). In all cases, gates were set for single cells (FSC-A/FSC-H and SSC-A/SSC-H), followed by a lymphocyte gate (FSC-SSC). CD8^+^ OT-I cells were identified as indicated in Fig 5A. The antibodies used were as follows, all purchased from BioLegend: CD8a-PerCP-Cy5.5 (53-6.7), Thy1.1-AF488 or -PE (OX-7), Thy1.2-BV510 (53-2.1), CD44-APC-Cy7 (IM7), CD62L-PE-Cy7 (MEL-14), IFNγ-PE (XMG1.2), IL-17A-PE-Cy7 (TC11-18H10), CD103-APC(2E7), CD69-PE(H1.2F3).

### Statistical Analysis

Statistical tests were performed as indicated in figure legends using GraphPad Prism 5.0.

### Ethics Statement

All animal experiments were carried out by following recommendations and approval from the Pennsylvania State University Animal Care and Use Committee (protocols 45613, 45794 & 46070) with great care taken to minimize suffering of animals.

## RESULTS

### Generation of *F.tularensis* LVS expressing OVA

We sought to constitutively express a fragment of the chicken ovalbumin protein, OVA_239-345_, in *F. tularensis* LVS in order to characterize the development and maintenance of antigen-specific CD8^+^ T cells in mice. A codon-optimized OVA_239-345_ tethered to a 6X- histidine tag at the C terminal end was ligated to the 3’ end of *vgrG* downstream of the highly active bacterioferritin promoter [29], generating pKK214-vgrG-OVA (Fig 1A). This was introduced into *F. tularensis* LVS, and expression determined by immunoblot of bacterial lysates using anti-6X histidine antibody. As shown in Fig 1B, a high level of expression of vgrG-OVA_239-345_ was observed.

**Figure 1:**
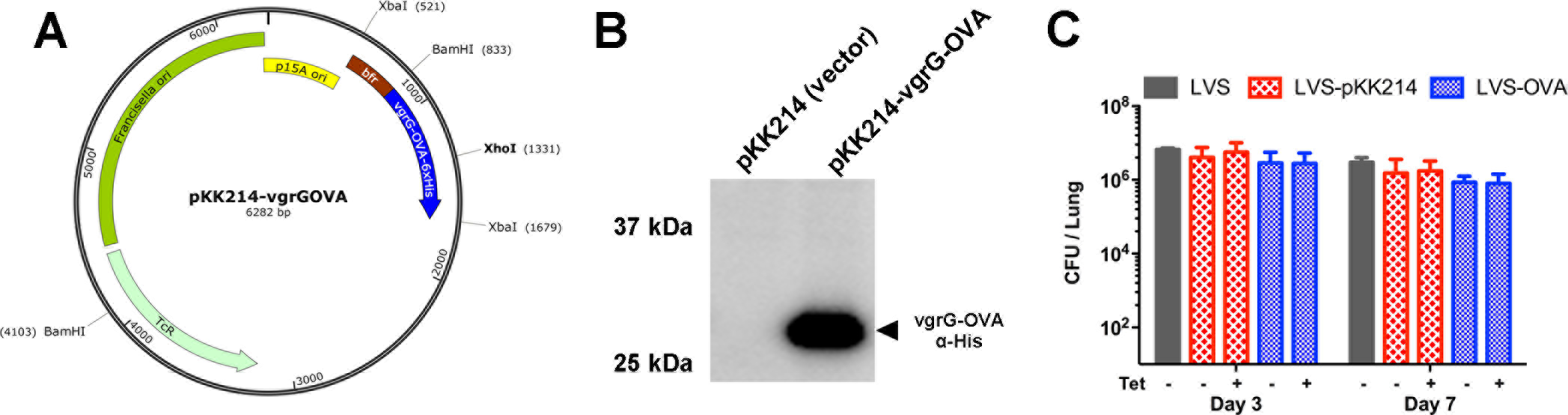
Generation and stability of LVS-OVA. A) Map of the pKK214-vgrG-OVA plasmid. B) Immunoblot demonstrating expression of the VgrG-OVA construct by cells harboring plasmid (pKK214-vgrG-OVA). C) Stability of the pKK214-vgrG-OVA plasmid during an *in vivo* infection. C57BL/6J mice (n=3 per group) were infected with 1000 CFU of wild-type *F. tularensis* LVS, LVS harboring empty vector (LVS-pKK214), or LVS harboring pKK214-vgrG-OVA (LVS-OVA). Serial dilutions of lung homogenates were plated with or without tetracycline as indicated.

Despite previous work showing that pKK214 plasmids are stably maintained without selection [30], we assessed the stability of pKK214-vgrG-OVA without selection i*n vivo*. Similar numbers of wild-type LVS, LVS-harboring empty vector pKK214, and LVS-OVA were recovered from the lungs of separate sets of mice on days 3 and 7 following infection with 1,000 cfu of each strain (Fig 1C). The presence or absence of tetracycline in culture plates did not affect the number of colonies of LVS-pKK214 or LVS-OVA recovered from infected mice (Fig 1C). These data show that our pKK214-vgrG-OVA expression construct allows LVS to express high levels of OVA and that the plasmid is robustly maintained without antibiotic selection *in vivo*.

### Antigen-specific proliferation of CD8^+^ OT-I cells in response to LVS-OVA

To test the ability of LVS-OVA to activate antigen-specific OT-I (CD8^+^) cells, we utilized an adoptive transfer model utilizing wild-type Thy1.1^+^ /Thy1.2^+^ congenic recipient mice. One day prior to infection with LVS-OVA, 5×10^6^ OT-I splenocytes (Thy1.1^−^/Thy1.2^+^) were transferred into wild-type congenic mice by intraperitoneal injection. Prior to injection, OT-I splenocytes were stained with CFSE, a bright and stable dye that is diluted with each round of cell division [31,32]. On days 5, 7, and 9 post-inoculation with LVS-pKK214 (vector control) or LVS-OVA, the lungs, lung-draining mediastinal lymph nodes (MLN), and spleens were collected and analyzed by flow cytometry. LVS-OVA led to robust proliferation of antigen-specific OT-I cells starting at approximately day 7 post-inoculation, while LVS-pKK214 was unable to stimulate such expansion (Fig 2A-C). By day 7 post-inoculation, many CFSE^+^ OT-I cells appeared to have undergone at least 5 rounds of cell division (Fig 2B). On day 9, OT-I cells represented the majority of the CD8^+^ T cells present in the lung (~60%) and a large proportion of the spleen (~25%), but were not as abundant in the MLN, suggesting that upon activation OT-I cells largely migrate out of the MLN (Fig 2C). Together, these data show that LVS-OVA specifically activate OT-I T cells and can be used to study CD8^+^ T cell responses to *Francisella*.

**Fig 2:**
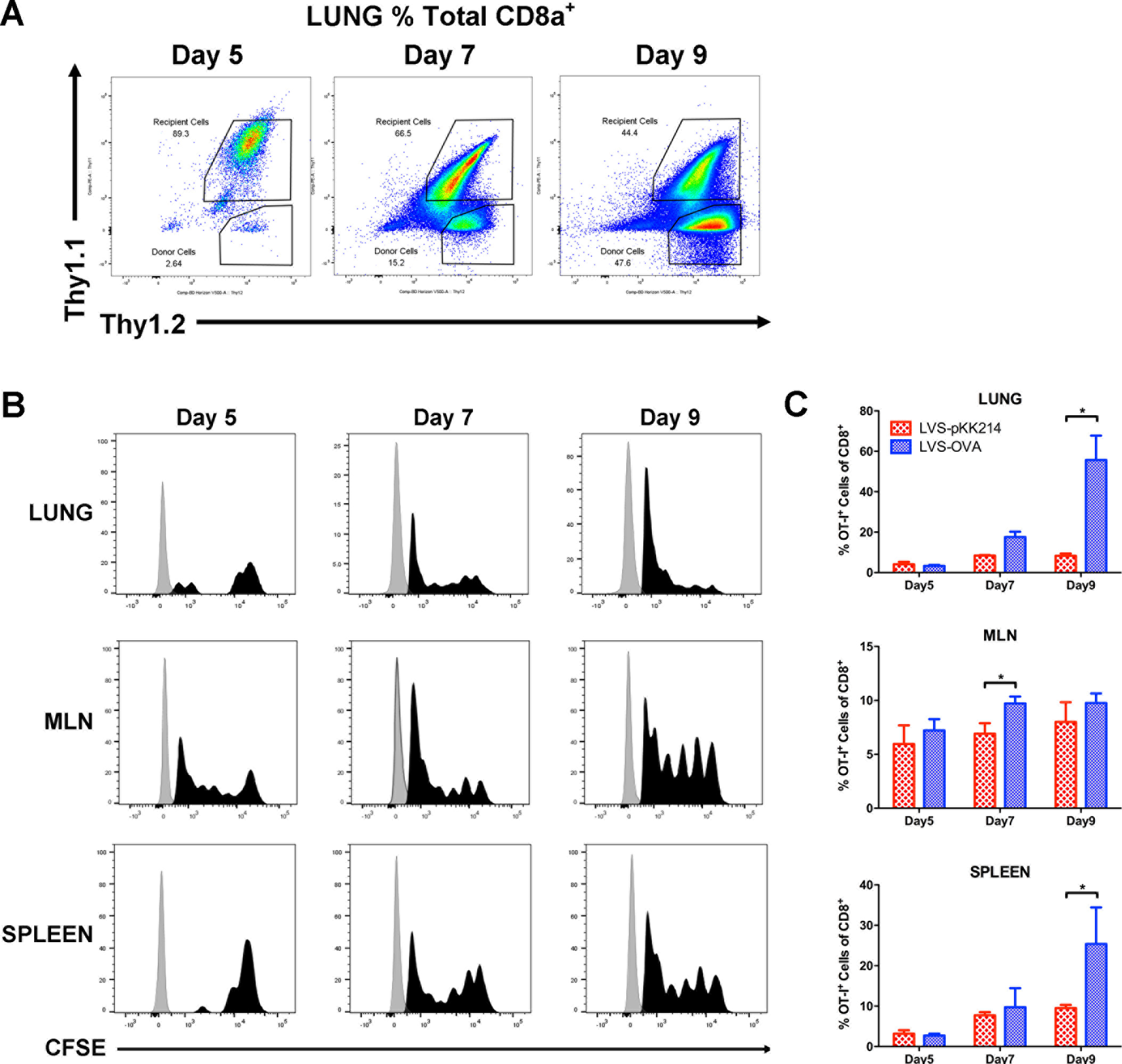
Antigen-specific proliferation of CD8^+^ OT-I cells in response to LVS-OVA. Thy1.1^−^ /Thy1.2^+^ OT-I cells were transferred in to Thy1.1^+^/Thy1.2^+^ recipient mice, 24 hours prior to infection with 1000 CFU of LVS-OVA or LVS-pKK214 (LVS with empty vector). A) representative scatterplots of CD8^+^ cells from the lungs of animals infected with LVS-OVA. B) CFSE dilution of CD8^+^ OT-I cells isolated from animals infected with LVS-OVA. C) The frequency of CD8^+^ OT-I cells in animals infected with LVS-OVA or LVS-pKK214. n=3 per group, means +/− SD are plotted; * p<0.05 according to two-tailed, unpaired T test.

### Kinetics of the CD8^+^ T cell response to intranasal LVS infection

To-date, it has not been possible to track the *Francisella*-specific CD8^+^ T cell response over time, due to a lack of tools to differentiate these cells from the broader CD8^+^ population. After adoptively transferring OT-I cells and infecting with LVS-OVA, we measured the number of OVA-specific CD8^+^ T cells in the lungs, MLN and spleen during infection and following bacterial clearance (Fig 3). Peak bacterial burdens were observed between day 3-7 post-infection, with far lower burdens evident on day 14, and clearance by day 21 (Fig3A). We observed a pattern similar to other pathogens where the T cell response peaks, then rapidly contracts after the infection is cleared. The scale and speed of contraction was greatest in the lungs; with a dramatic reduction in the number of OT-I cells from day 7 to all later time points (Fig 3B). In the MLN and spleen, contraction was more gradual, with retention of a greater number of OT-I cells through day 71 (Fig 3C-D). These data suggest that *Francisella*-specific T cells are retained poorly in the lungs compared to secondary lymphoid organs, even following intranasal vaccination.

**Fig 3:**
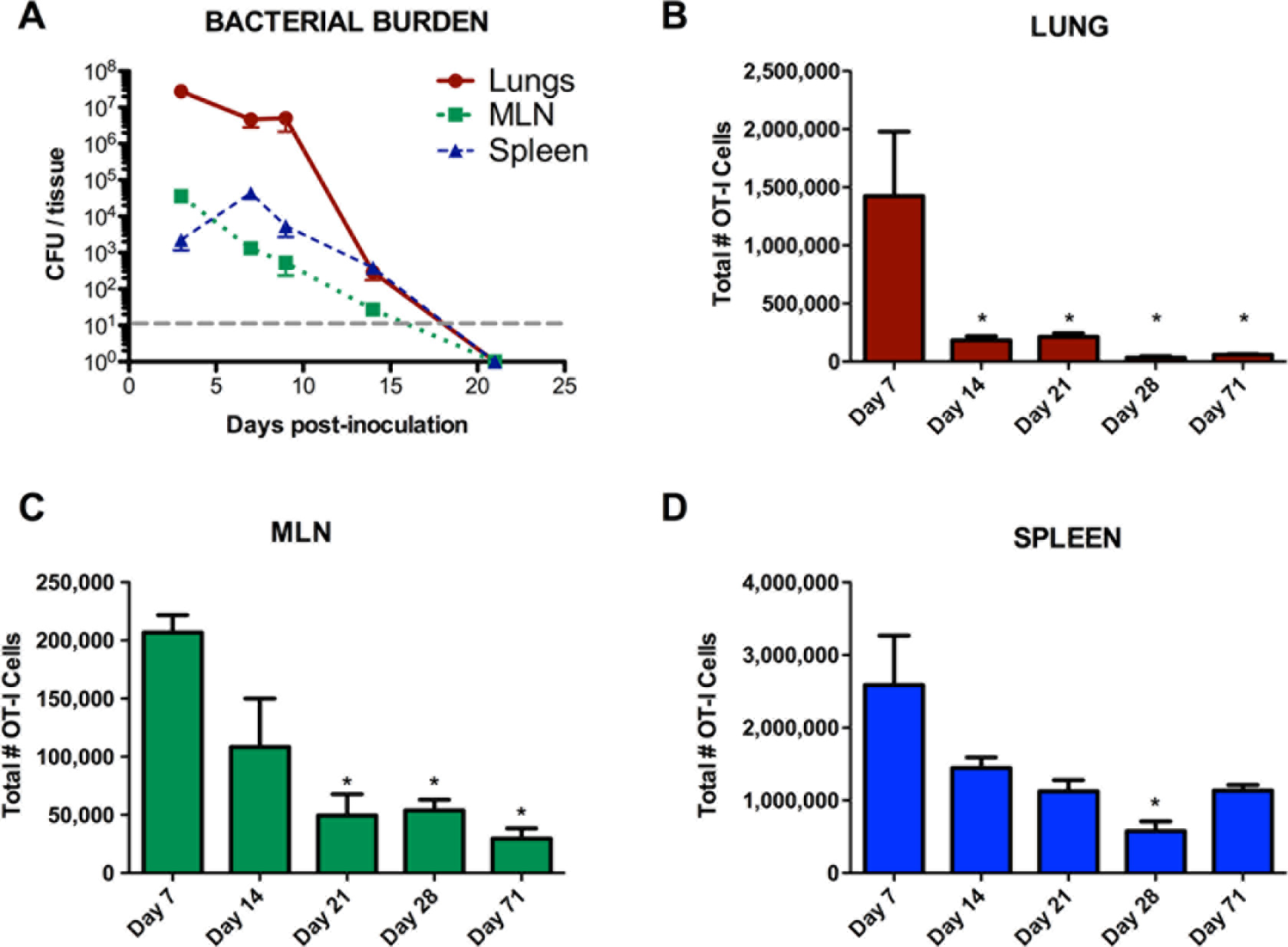
Kinetics of the CD8^+^ T cell response to intranasal LVS infection. OT-I cells were adoptively transferred into recipient mice 24 hours prior to infection with 1000 CFU of LVS-OVA. A) Bacterial burden of infected animals over time. B-D) number of CD8^+^ OT-I cells in infected animals over time. n=3 per group, means +/− SEM are plotted. Statistical significance (p<0.05) for one-way ANOVA with Tukey’s post test, for the indicated column compared to: * day7.

### *Francisella*-specific CD8 IFNγ and IL-17A Responses

One of the primary effector functions of CD8^+^ T cells is the production of inflammatory cytokines. Activated CD8^+^ T cells are capable of producing a range of cytokines depending on inflammatory cues and transcriptional programing [33,34]. Among these, IFN-γ is known to play an important role in activating phagocytic cells to kill many intracellular pathogens, including *F. tularensis* [35–37]. The role of IL-17A in the immune response to *F. tularensis* has been studied, but has been more difficult to define, and appears to vary based on *F. tularensis* strain, route of exposure, and primary versus recall response [37–40]. To study the production of these two representative cytokines by *Francisella*-specific CD8^+^ T cells, we performed intracellular cytokine staining on cells isolated from LVS-OVA-infected animals and stimulated with PMA/calcium ionophore.

As expected, we observed that some CD8^+^ OT-I cells produced IFN-γ during and after infection with LVS-OVA (Fig 4A). For each organ, the peak in IFN-γ producing cells coincided with peak T_EM_ responses (see below), and occurs around the time the immune response is beginning to contain bacterial burden. We did not observe any significant production of IL-17A by stimulated OT-I cells at any time-point during infection (Fig 4B). These results show that infection with LVS via the intranasal route elicits *Francisella*-specific CD8^+^ T cells that produce IFN-γ but not IL-17A.

**Fig 4:**
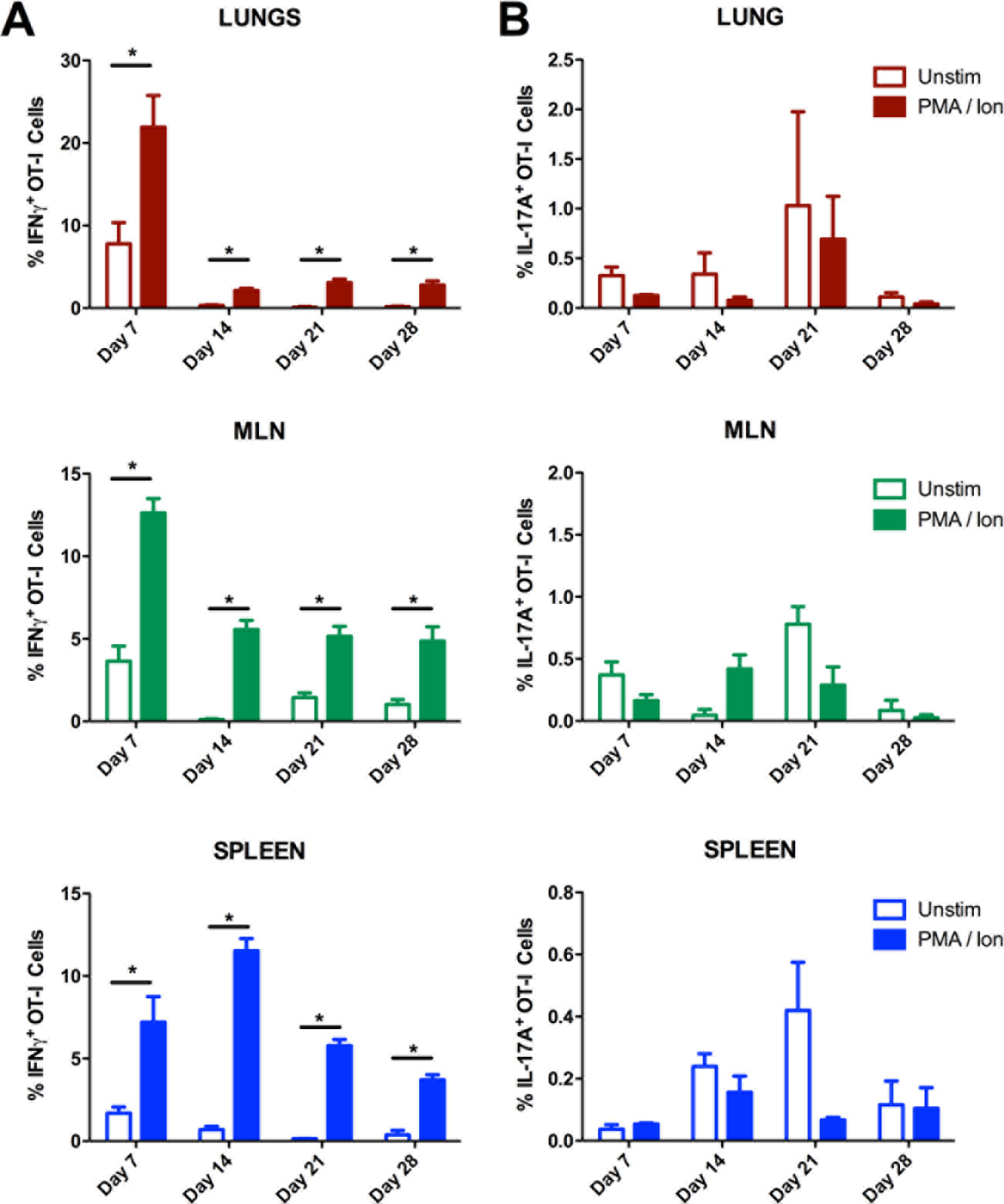
*F. tularensis*-specific CD8^+^ IFN-γ and IL-17A responses. OT-I cells were adoptively transferred into recipient mice 24 hours prior to infection with 1000 CFU of LVS-OVA. Intracellular cytokine staining was performed on single cell suspensions incubated with BrefeldinA, +/− PMA and calcium ionophore. n=3 per group, means +/− SEM are plotted. * p<0.05 according to two-tailed, unpaired T test.

### *Francisella*-specific CD8^+^ memory T cell development

Upon activation, naïve CD8^+^ T cells differentiate into effector and memory subsets. Effector/memory (T_EM_) CD8^+^ T cells are characterized by the decreased expression of markers associated with lymphoid tissue homing (eg: CD62L) and increased expression of markers associated with homing to the source of inflammation (eg: CD44) [41]. These cells have a high propensity for effector functions, including killing infected cells and producing cytokines.
Central memory (T_CM_) cells are less geared towards effector functions; rather they retain extensive proliferative capacity, and primarily circulate through the secondary lymphoid organs due to high expression of CD62L and other lymphoid tissue homing markers [41]. The primary role of T_CM_ is generally considered to be their ability to rapidly expand into a new population of effector cells upon re-exposure to antigen [42,43]. Using our model, we monitored the *Francisella*-specific T_EM_ and T_CM_ subsets that arose in the lungs, MLN and spleen in response to intranasal LVS-OVA infection.

In the lungs, the highest number of effector memory OT-I cells were observed on day 7 (Fig 5B), which coincides with the peak bacterial burden in this assay (Fig 3A). T_EM_ remained at high levels in the lung to the time of clearance then dropped sharply in number, with only a small number detected on d28 post-infection. In the MLN, T_EM_ numbers followed a similar general pattern as in the lung, with the most cells present on day 7; however, a transient increase was observed on day 28, possibly reflecting egress of these cells out of the lung (Fig 5B). In contrast to the lungs and MLN, T_EM_ peaked in number on day 14 in the spleen (Fig 5B). This may be due to the fact that bacteria reach the spleen and replicate later in the course of pneumonic tularemia compared to the lungs (Fig 3A and [44,45]). Consistent with the data for total OT-I cells (Fig 3), *Francisella*-specific T_EM_ are retained well in the spleen and poorly in the lungs after bacterial clearance.

**Fig 5:**
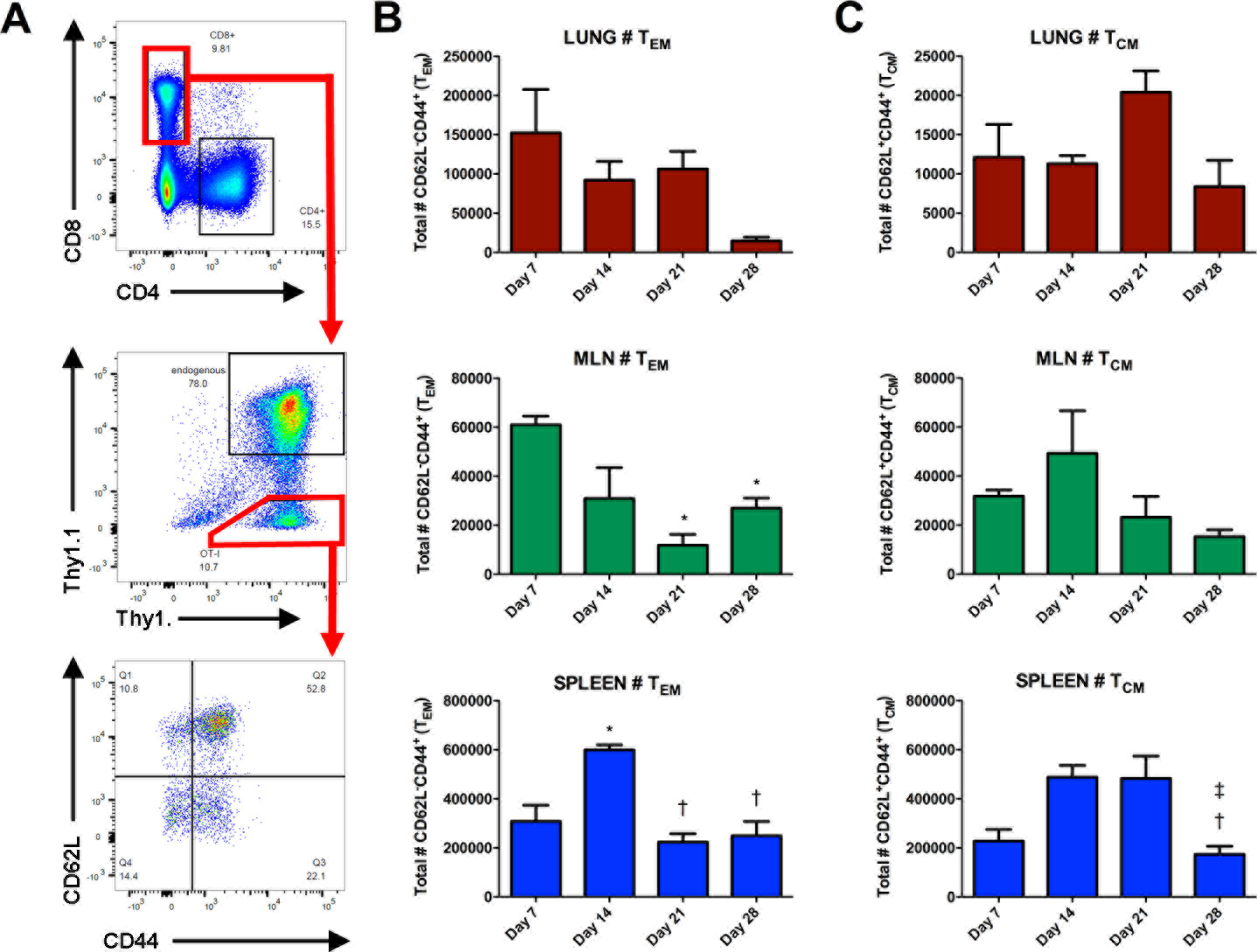
*F. tularensis*-specific CD8^+^ memory T cell development. OT-I cells were adoptively transferred into recipient mice 24 hours prior to infection with 1000 CFU of LVS-OVA. A) Representative scatterplots demonstrating gating strategy. B) number of T_EM_ OT-I cells. C) number of T_CM_ OT-I cells. n=3 per group, means +/− SEM are plotted. Statistical significance (p<0.05) for one-way ANOVA with Tukey’s post test, for the indicated column compared to : * day7, † day 14, ‡ day 21.

As expected, the number of T_CM_ in the lung was much lower than the number of T_EM_ during infection, and remained similar from day 7 to day 28 (Fig 5C). The number of T_CM_ in the MLN peaked on day 14, then exhibited a trend towards gradual decrease over time. In the spleen, T_CM_ increased in number between day 7 and 21, and decreased on day 28 (Fig 5C). These data show that our LVS-OVA/OT-I model can be used to monitor *Francisella*-specific CD8^+^ T cell memory subsets, and that these subsets show disparate expansion, contraction and retention properties in various tissues following intranasal LVS infection.

### Comparison of *Francisella*-specific CD8^+^ T cell responses to scarification-prime, intranasal boost-versus intranasal-prime, intranasal-boost vaccination strategies

To demonstrate one potential application of LVS-OVA, we studied CD8^+^ T cell responses in mice primed either intranasally or via scarification, both followed by an intranasal booster. Responses were assessed on day 28 after primary vaccination, and also three and five days after an intranasal booster given on day 30. Mice primed by the intranasal route exhibited significant weight loss before recovering, while mice primed via scarification did not lose weight (Fig 6A). The intranasal booster was well tolerated by both groups (Fig 6B and Fig S1). On day 28 post-vaccination, similar numbers of OT-I cells were present in the lungs and spleens of both groups (Fig 6C). The MLNs of animals given the intranasal primary vaccination harbored more OT-I cells at this point (Fig 6C). The intranasal booster increased the number of OT-I cells in the MLNs of scarification-primed mice, such that they matched intranasally-primed mice. Additionally, the number of *Francisella*-specific cells in the lungs of scarification-primed mice increased after the booster, reaching significantly higher levels than in intranasally-primed and boosted mice. A similar trend was observed for the numbers of *Francisella*-specific T_EM_ and T_CM_ cells following priming and booster in both schemes of vaccination (Fig 6D-E). Finally, we assessed the number of OT-I cells displaying the resident memory markers CD69 and CD103 within the lungs after these vaccination schemes (Fig 7). For all animals, very few CD69^+^ CD103^+^ cells were detected in the lungs, possibly reflecting the poor development of resident memory cells in this model. Altogether, these data suggest that: 1) Intranasal LVS vaccination leads to more *Francisella*-specific CD8^+^ T cells in the MLN compared to scarification vaccination; 2) an intranasal booster increases the number of *Francisella*-specific CD8^+^ cells in the MLN and lung of scarification-primed animals, while avoiding the weight loss associated with intranasal primary vaccination; and 3) neither regimen appears to effectively elicit *Francisella*-specific lung-resident CD8^+^ T cells.

**Fig 6:**
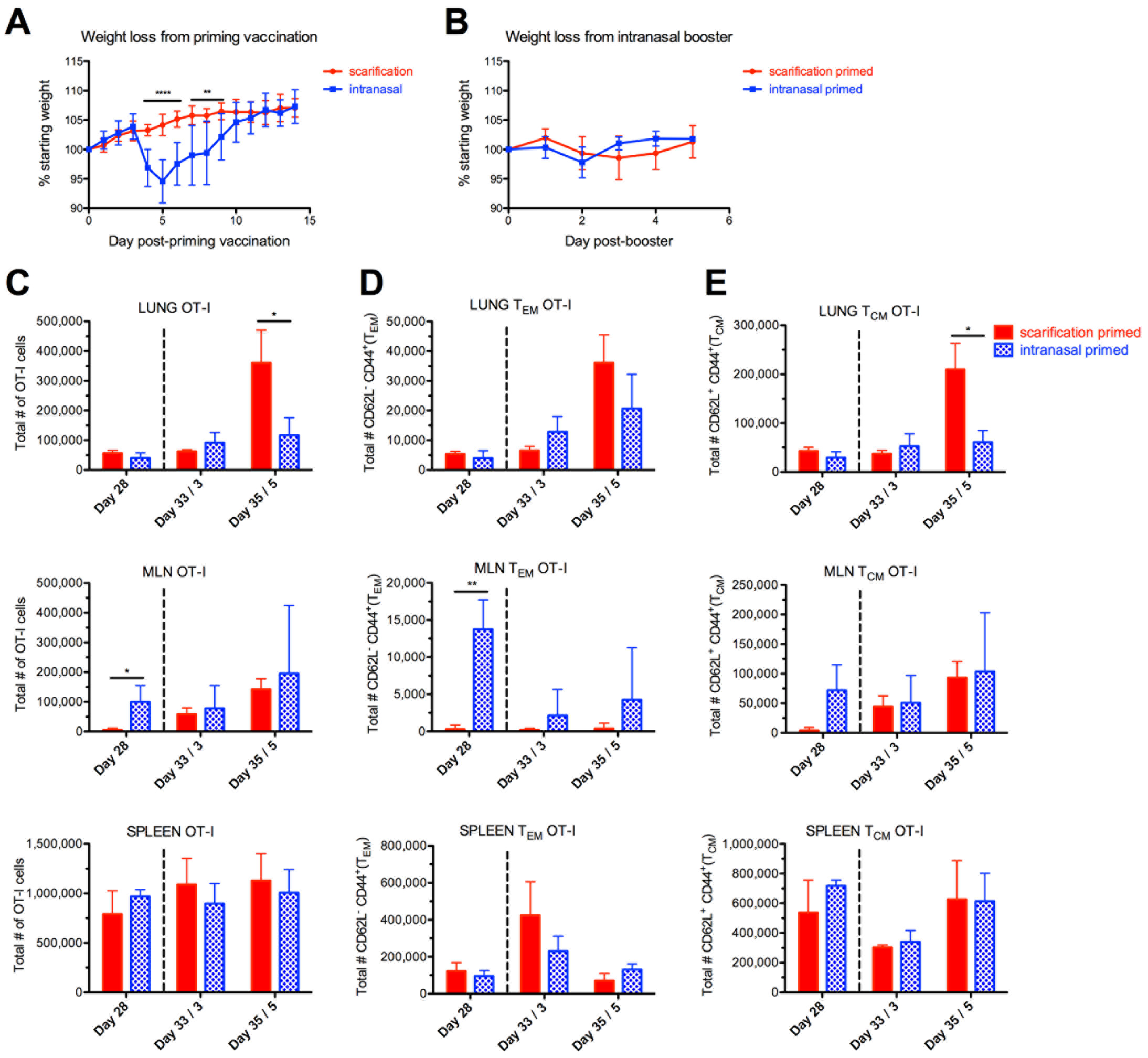
Comparison of *F. tularensis*-specific CD8^+^ T cell responses from scarification-prime intranasal-boost, versus intranasal-prime intranasal-boost vaccination strategies. OT-I cells were adoptively transferred into recipient mice, 24 hours prior to priming vaccination with LVS-OVA via the intranasal or scarification route. An intranasal booster was given on day 30. A) weight loss of mice from the priming vaccination. **p<0.01 and ***p<0.001, two-tailed unpaired T test. B) weight of mice following intranasal booster. C-E) The total numbers of OT-I cells (C), T_EM_ OT-I cells (D) and T_CM_ OT-I cells (E) were determined 28 days after priming vaccination, and three and five days after booster (represented by dotted line). Statistical significance was determined by two-tailed, unpaired t-test at each timepoint * p<0.05 ** p<0.01. n=3 per group, means +/− SD are plotted.

**Fig 7:**
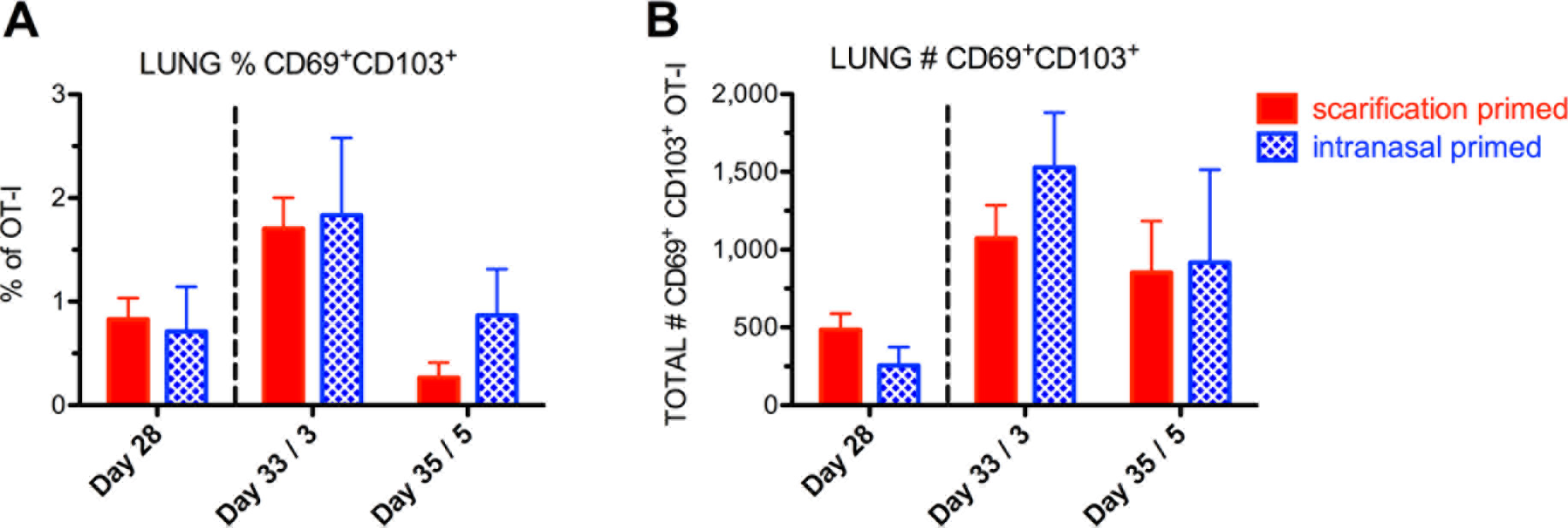
*F. tularensis*-specific lung CD8^+^ T cells with resident memory markers following prime-boost vaccination strategies. A) relative frequency, and B) number of CD69^+^CD103^+^ OT-I cells. n=3 per group, means +/− SD are plotted. No statistically significant differences according to two-tailed, unpaired t-tests.

## DISCUSSION

There is currently no approved vaccine for the prevention of tularemia, due to concerns regarding the safety and efficacy of existing vaccine candidates. Progress towards a vaccine has been hampered by a lack of understanding of the correlates of protective cell-mediated immunity, in part due to a lack of tools to identify these correlates. Here we report the development of an ovalumbin expression system for *F. tularensis,* which is the first tool to allow for the study of *F. tularensis*-specific CD8^+^ T cells. Our early attempts to express ovalbumin in *F. tularensis* were unsuccessful. However, after codon optimizing a fragment of ovalbumin, and expressing it as a C-terminal tag of the native VgrG protein did we achieve a robust expression (Fig 1A-B). A strain of *F. tularensis* LVS containing our expression construct (LVS-OVA) stably maintained the plasmid without selective antibiotics, and stimulated the expansion of adoptively-transferred CD8^+^ OT-I cells in infected mice.

We first sought to study the initial expansion of *F. tularensis*-specific CD8^+^ T cells after intranasal infection. Expansion primarily occurred between days 7-9, with relatively little division of OT-I cells evident on day 5. This contrasts with the kinetics observed for another cytosolic bacterial pathogen, *Listeria monocytogenes,* which elicits significant expansion of CD8^+^ T cells by day 3 [46]. *F. tularensis* employs a stealth strategy that delays activation of innate pro-inflammatory pathways early in the course of infection [10,44,45,47]; these data suggest this stealth strategy may also dramatically affect the kinetics of the T cell response. Our data suggest the peak of the CD8^+^ T cell response to *F. tularensis* is likely somewhere between day 9 and 14 (Fig 2-3). During infection, CD8^+^ T cells help to eliminate pathogens by killing target cells and producing inflammatory cytokines. Unsurprisingly, we observed that *F. tularensis*-specific CD8^+^ T cells produced IFN-γ upon stimulation (Fig 4). We did not observe IL-17 producing OT-I cells at any point. In the future, LVS-OVA could be used to determine what other effector molecules are produced by *F. tularensis*-specific CD8^+^ T cells, and whether effector profile is altered by other factors, such as vaccine strain or route of exposure.

A major advantage of LVS-OVA is that it allows us to study pathogen-specific memory development, particularly at times when cells of other specificities may be more abundant. We have observed that *F. tularensis*-specific CD8^+^ T_CM_ and T_EM_ subsets show disparate expansion, contraction and retention properties in various tissues following intranasal LVS infection (Fig 5). Notably, peaks in the T_EM_ population in each organ coincide with peaks in IFN-γ producing cells, and occur around the time bacterial burden is beginning to decline. After clearance of primary infection, T_EM_ are poorly retained in the lungs. The recently described resident memory subset of T cells (T_RM_) remain in a tissue after the clearance of primary infection, and are thought to provide a rapid response that contributes to protection from reinfection, including against respiratory pathogens [48,49]. However, we detected very few *F. tularensis*-specific CD8^+^ T cells expressing the resident memory markers CD69 and CD103 in the lungs of vaccinated mice. This suggests that lung T_RM_ development is poor in this model, and that strategies to boost *F. tularensis*-specific T_RM_ in the lungs may enhance the protectiveness of vaccines. Notably, the generation of lung T_RM_ has been associated with tissue regeneration after injury [49], and the RML strain of LVS, which confers superior protection against virulent strains, has also been shown to be more virulent [10]. It is interesting to speculate that the increased protectiveness of the RML LVS strain may be due to increased generation of *F. tularensis*-specific T_RM_ in the lungs. Studies using our ovalbumin expression system to compare effective and ineffective vaccine responses will help to define correlates protective adaptive immunity.

An ideal vaccine would preferentially induce such protective responses without causing any harm to the host. In this regard, vaccination with *F. tularensis* LVS by the respiratory route appears to provide superior protection compared to other routes [12,19,50,51], but is also associated with significant adverse effects [50]. We used LVS-OVA to compare the CD8^+^ T cell responses elicited by intranasal vaccination versus skin scarification, a more commonly used route in humans. Mice vaccinated via the intranasal route exhibited significant body weight loss before recovering, while mice vaccinated via scarification did not lose weight (Fig 6). Intranasal vaccination led to more *F. tularensis*-specific CD8^+^ T cells in the lung-draining MLN; however, after an intranasal booster this difference was no longer evident. The intranasal booster also increased the number of *F. tularensis-specific* CD8^+^ T cells in the lungs of scarification-primed animals, but they did not lose any weight and tolerated the booster similarly to intranasal-primed animals. One possible implication of these data is that a scarification-prime, respiratory-boost strategy may achieve the same benefits as respiratory vaccination, with enhanced safety.

In summary, we have developed an ovalbumin expression system that allows for the identification and study of *F. tularensis*-specific CD8^+^ T cells. Using an adoptive transfer model, we demonstrate that this tool can be used to study these cells’ expansion, effector functions, retention as memory cells, and response to re-challenge or booster vaccinations. The availability of this system opens up a number of possibilities for critical detailed studies, including how CD8^+^ T cell responses vary based on *F. tularensis* strain, route of inoculation, available cytokines, and how adjuvants may enhance these responses. Ultimately, such studies will enhance our understanding of protective adaptive immunity, and helping to lay the foundation for the development of a safe and effective vaccine for the prevention of tularemia.

## ACKNOWLEDGMENTS

We sincerely thank Dr. Karl Klose, University of Texas San Antonio for providing plasmid pKK214. We also thank The Huck Institutes of the Life Sciences for the facilities and technical support.

## SUPPORTING INFORMATION CAPTIONS

**Fig S1:**
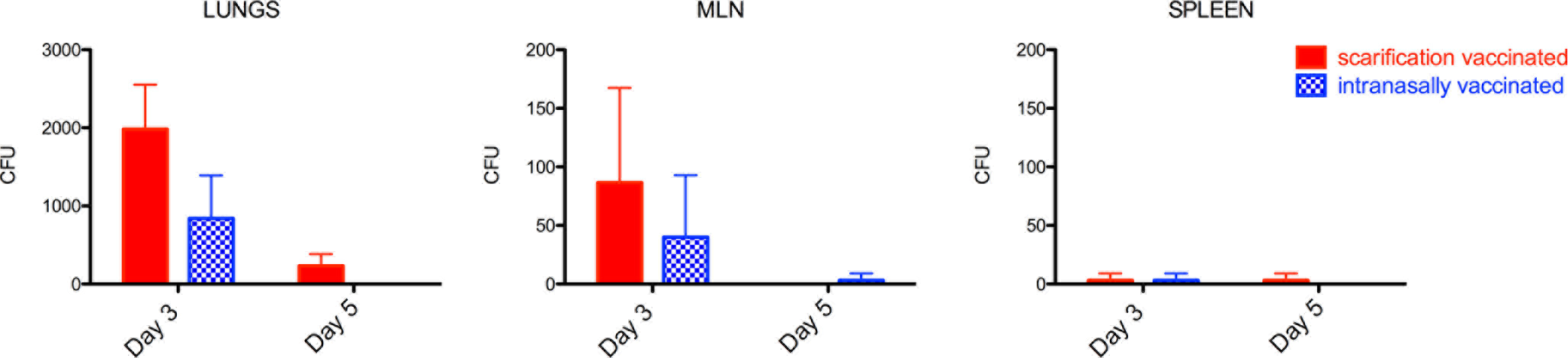
Bacterial burdens following intranasal booster. Mice (n=3 per group) were given a priming vaccination of LVS-OVA via the scarification or intranasal route, followed by a booster with 10,000 CFU on day 30. Bacterial burdens are shown on day 3 and day 5 after the booster.

